# Novel oil-associated bacteria in Arctic seawater exposed to different nutrient biostimulation regimes

**DOI:** 10.1101/2024.03.22.586348

**Authors:** Francisco D. Davila Aleman, María A. Bautista, Janine McCalder, Kaiden Jobin, Sean M. C. Murphy, Brent Else, Casey R.J. Hubert

**Affiliations:** Department of Biological Sciences, University of Calgary, Calgary, Alberta, Canada; Department of Geography, University of Calgary, Calgary, Alberta, Canada

**Keywords:** Microcosms, Arctic, Amplicon sequencing, oil-degraders, biostimulation

## Abstract

The Arctic Ocean is an oligotrophic ecosystem facing escalating threats of oil spills as ship traffic increases owing to climate change-induced sea ice retreat. Biostimulation is an oil spill mitigation strategy that involves introducing bioavailable nutrients to enhance crude oil biodegradation by endemic oil-degrading microbes. For bioremediation to offer a viable response for future oil spill mitigation in extreme Arctic conditions, a better understanding of the effects of nutrient addition on Arctic marine microorganisms is needed. Comprehensive population tracking of controlled oil-spill microcosms using cell counting and microbial biodiversity screening revealed a significant decline in community diversity together with changes in microbial community composition. These shifts were also indicative of changes in prevailing genomic traits as inferred from 16S rRNA taxonomy of resulting communities. In addition to well-recognized hydrocarbonoclastic bacteria, differential abundance analysis highlighted significant enrichment of unexpected genera *Lacinutrix*, *Halarcobacter* and Candidatus *Pseudothioglobus*. These groups have not been associated with hydrocarbon biodegradation until now, even though genomes from closely related isolates confirm the potential for hydrocarbon metabolism. These findings broaden understanding of marine oil spill bioremediation and how Arctic marine microbiomes and their novel lineages can respond to nutrient biostimulation as a strategy for oil spill mitigation.

**Importance:** A comprehensive characterization and understanding of the impact of marine bioremediation strategies in the Arctic is crucial for effectively managing oil contamination. Such understanding enables an evaluation of the ecological impacts associated with mitigation strategies to minimize negative effects on sensitive ecosystems. By introducing external nutrients into areas affected by spills, microbial growth can be stimulated, enhancing hydrocarbon degradation by naturally occurring oil-degrading microorganisms. This may include novel microbial groups in permanently cold Arctic waters, where fewer oil biodegradation studies have been performed. It is also important to consider how nutrient addition may affect endemic microbial communities after successful remediation has occurred in oil-contaminated zones. Promoting naturally occurring oil-degrading microorganisms may have significant implications on nutrient cycling and marine food webs, which are critical for sustaining the health and well-being of coastal Indigenous communities in northern latitudes.

## Introduction

Anthropogenic activity has rapidly increased greenhouse gas emissions contributing to a significant rise in temperatures in polar regions (1). Melting sea ice and glaciers in the Arctic are facilitating increased maritime navigation and elevated risks of pollution events such as spills of crude oil, bunker fuel or diesel (2). The Arctic marine environment is inhabited by microbial populations that contribute significantly to the biogeochemical cycling of essential macro- and micronutrients that are vital to the growth and sustenance of higher trophic levels (3). Microbiomes thus form the base of complex oceanic food webs (4). Understanding how microbial life in the Arctic Ocean will respond to anticipated environmental perturbations such as accidental oil spill events requires careful investigation.

The Arctic Ocean is geographically unique, encircled by land with only two main connections to the Pacific and Atlantic oceans. Rivers draining into the Arctic supply relatively more freshwater than any other ocean basin receives, but this tends to be constrained to a narrow riverine coastal domain (5). These dynamics contribute to much of the Arctic Ocean behaving as an oligotrophic ecosystem characterized by low levels of nitrogen and phosphorus (6), restricting microbial growth and metabolism (7, 8). Accordingly, biodegradation of spilled oil by naturally occurring microbial populations will also be nutrient-limited. To address this limitation and enhance the breakdown of crude oil, biostimulation can be considered as an oil spill mitigation strategy in the Arctic marine environment. Bioavailable nutrients, such as N- and P-containing compounds, can promote biodegradation of crude oil by alleviating nutrient limitation on oil-degrading microbial populations within resident microbiomes (9, 10). While bioremediation through nutrient addition stands as a viable solution to respond to oil spills, the introduction of nutrients into the environment can in its own right be considered an ecological disturbance (11), i.e., altering an environment’s taxonomic composition or physical properties (10, 11). Introducing exogenous surfactants or nutrients can increase rates of hydrocarbon biodegradation and significantly shift microbial community composition, enriching hydrocarbonoclastic bacterial genera (e.g., *Colwellia*, *Oleispira*, *Alcanivorax*, *Marinobacter*, *Pseudomonas* and *Cycloclasticus*) and decreasing microbial community diversity (10, 12–14). A recent study highlighted that this response can be variable in the marine environment based on factors such as geographic location and temperature determining differences in microbial activity, diversity, community composition and growth rate (15).

Microbial interactions in the ocean are vital for maintaining ocean ecosystem function. Abiotic factors such as nutrient availability strongly affect these interactions, leading to a higher rate of positive bacterial associations under oligotrophic environments compared to high nutrient environments that feature greater competition between microorganisms (16). The extent to which these effects hold true in the context of nutrient biostimulation for oil biodegradation in the cold waters of the Arctic Ocean remain relatively unexplored (17). Beyond shifts in microbial diversity, universal genomic traits such as genome size, GC content and number of coding sequences are important parameters related to how microbiomes respond to nutrient availability (16, 18–21). Larger genome size has been correlated with high nutrient environments compared to unenriched ecosystems, indicating a community adaptation connected with genetically encoded phenotypes (18).

The present study investigates how an Arctic marine microbial community responds to different nutrient levels by examining microbial growth, hydrocarbon biodegradation rates, changes in microbial community composition, inter-species co-occurrence and inferred genomic traits. The study was conducted using seawater collected in Canada’s Kitikmeot Sea (69°N), an enclosed basin with significant river input (22). This oligotrophic system serves as a useful analog for the Arctic Ocean more generally with very low N and P concentrations (23) and seasonal sea ice cover (24). Mock oil spill microcosms enabled testing the hypothesis that oil-contaminated systems subjected to nutrient addition result in the enrichment of hydrocarbonoclastic bacteria characterized by larger genome size and higher GC content, concomitant with increased rates of hydrocarbon degradation.

## Materials and Methods

### Sample collection

Seawater was collected in May 2019 close to the community of Iqaluktuuttiaq (Cambridge Bay), Nunavut (69°06′29″N, 105°03′33″W). Seawater was sampled from directly underneath 1.7 meter thick sea-ice by drilling a hole through using a 25 cm diameter auger. A peristaltic pump was used to collect 20 L of under-ice seawater, which was transported by snowmobile to the Canadian High Arctic Research Station in Iqaluktuuttiaq. Accordingly, seawater samples were able to be processed within 8 hours of collection.

### Media and microcosm establishment

To assess the impact of nitrogen and phosphorous availability on hydrocarbon biodegradation by the microbial community in the under-ice seawater, three different culture media conditions were implemented. For high nutrient and low nutrient conditions, artificial seawater medium ONR7a (25) and dehydrated sea salt crystals (Red Sea Salt medium; Red Sea Fish Pharm LTD) were used, respectively. Relative to the seawater in this region, levels of nitrogen and phosphorous are approximately 10 times greater in high-nutrient ORN7a medium (23), which is a standard growth medium for cultivating hydrocarbonoclastic marine bacteria under laboratory conditions (23, 25–27). Additionally, seawater from the sampling location was double-pasteurized (80°C for 1h) and used as a third medium to represent ambient nutrient levels at the sampling site. Replicate microcosms were established in sterile 160 ml glass serum bottles by combining 25 mL of under-ice seawater as an inoculum with 25 mL of the different medium types, leaving 110 ml of headspace. Six microcosm bottles per condition were amended with light crude oil to a final concentration of 0.1% v/v. Subsequently, bottles were capped with autoclaved 20 mm butyl rubber stoppers and aluminium caps. The crude oil used comes from the Macondo site in the Gulf of Mexico and is a light, non-viscous oil with an initial API gravity of 35° (12). Three additional microcosm bottles per medium condition received no crude oil amendment in order to test the effect of nutrient addition on the microbiome without crude oil present. Microcosms were placed in a shaking incubator at 140 rpm at 4°C in the dark for 15 weeks. Subsampling involved 10 mL aliquots being removed after 5, 10 and 15 weeks to facilitate DNA extraction (9.5 mL) and cell counting (0.5 mL) using a sterile syringe. Of the oil-amended microcosms for each medium type, one was sacrificed at the inoculation time point (T0), one after 5 weeks (T1), and one after 15 weeks (T3) to assess biodegradation through oil geochemistry analysis. All available replicate microcosm bottles were used for DNA extraction and cell counting (12) throughout the incubation period at T0 (7 replicates), T1 (6 replicates), T2 (5 replicates) and T3 (5 replicates).

### Fluorescence microscopy cell counting

To monitor microbial growth, fluorescence cell counting was performed as described elsewhere (28). Briefly, 0.5 mL was subsampled from each microcosm at different time points and fixed with 600 µl of 6% paraformaldehyde with addition of PBS to a final volume of 10 mL. Samples were incubated at room temperature for 1 h, followed by vacuum filtration (0.2 μm pore size, white polycarbonate, 47 mm diameter; Millipore, Eschborn, Germany). For DAPI staining, filters were sectioned into quarters, immersed in 10 mL of DAPI solution (1 µg/mL), and incubated at room temperature for 10 minutes in the dark. Filter quarters were briefly rinsed with MilliQ water and 80% ethanol, followed by air drying on Whatman paper. Finally, filter quarters were placed on glass slides with 10 µL of a mixture of vectashield and mounting medium (4:1) and counted using epifluorescence microscopy. Counting used the criteria previously described by Kirchman et al. (29). Three filter quarters were used per sample and 40 fields of view were counted per filter quarter. Each field and quarter were averaged to calculate the total number of cells per sample at each time point.

### Hydrocarbon extraction and quantification

Hydrocarbons from one crude oil-contaminated microcosm per medium condition were extracted at 0, 5 and 15 weeks using a previously reported protocol (12) involving dissolution in dichloromethane (DCM) and passage through a sodium sulphate column to remove residual water. After DCM evaporation, samples were reconstituted in 1 mL of DCM. Hydrocarbon extracts were injected for GC-MS under the same parameters previously reported (12). Ratios of nC_17_/pristane, nC_18_/phytane and 3-methylphenanthrene/9-methylphenanthrene (3MP/9MP) were determined and used as proxies for crude oil biodegradation on the basis that pristane, phytane and 9-methylphenanthrene are less susceptible to biodegradation than nC_17_, nC_18_ and 3-methylphenanthrene (30, 31).

### DNA extraction and sequencing

DNA was extracted using the DNeasy PowerWater Kit (QIAGEN) following manufacturer’s instructions with a few modifications. Bead beating was set to 2.4 m/s for 5 min in an Omni Bead Ruptor 24 (Omni International Inc). Samples were eluted in 50 µl of nuclease-free water pre-warmed to 50°C. DNA was quantified using a HS dsDNA Qubit (v.2.0) assay following manufacturer’s instructions and diluted to 5 ng/ul if the concentration was above this before proceeding to PCR. PCR amplicon libraries were generated for the V4 region of bacterial and archaeal 16 rRNA genes using primers 515FB (5’-GTGYCAGCMGCCGCGGTAA-3’) and 806RB (5’-GGACTACNVGGGTWTCTAAT-3’), as described previously (12, 32, 33). PCR was performed in 25 µl reaction volumes in triplicate for each sample using 2x KAPA HiFi HotStart ReadyMix. PCR products for each sample were pooled before bead cleanup, adaptor ligation and indexing. Amplicon libraries were quantified using a HS dsDNA Qubit (v.2.0) assay following manufacturer instructions. Libraries were sequenced using an in-house Illumina MiSeq and v2 reagent kit producing 2× 250 bp paired-end reads.

### Amplicon sequencing processing and diversity analysis

Raw sequencing data from paired-end reads were demultiplexed and exported as fastq files. DADA2 version 1.24.0 (34) with default parameters was used to generate amplicon sequence variants (ASVs) and detect and remove chimeras. ASVs were taxonomically classified using the most recent DADA2-formatted 16S rRNA database from GTDB-Tk (35) version 4.3 (released June 2022). ASV sequences were normalized using the cumulative sum scaling method. Alpha diversity (observed species) and beta diversity (Bray-Curtis dissimilarity) were quantified using the R package “microbiome” (36) version 1.18.0 and the R package “phyloseq” (37) version 1.48.0, respectively. Differences in microbial diversity were determined using Kruskal-Wallis tests using the R package ggpubr version 0.6.0. ASV relative abundances were estimated using the R package “microbiomeutilities” (38) version 1.00.16.

### Differential abundance analysis

Total number of 16S rRNA genes for each taxon were calculated by multiplying the relative abundance of each ASV in a given microcosm by the average cell count for that microcosm bottle. Rare ASVs were filtered out using a prevalence threshold <10% for each nutrient condition. Significant changes were analyzed using White’s non-parametric test with BH correction, 1000 permutations and visualized using the R package “microbiomeMarker” (39) version 1.3.3.

### Inference of community-weighted genomic traits

For inference of community-weighted genomic traits (genome size, GC content and CDS) the R package “paprica” (19) version 0.72 was used. Paprica implements a phylogenetic placement approach where sample amplicon reads (ASVs) are placed on a reference phylogenetic tree created from 16S and 23S rRNA genes from sequenced bacterial and archaeal genomes (19). Since the genomic traits of each phylogenetic edge of the reference tree are known, paprica generates an estimate of genomic traits for each of the ASVs placed on the reference tree (19, 40–43).

### Estimation of microbial ecological association

To infer microbial ecological associations, Sparse Inverse Covariance Estimation for Ecological Association Inference (SPIEC-EASI) with neighborhood selection method (MB) was implemented among crude-oil amended ASVs under the three distinct nutrient regimes using the R package “SpiecEasi” (44) version 1.1.1. For visualization of the inferred microbial networks, the R package “igraph” (45) version 1.3.4 was implemented. For calculating possible microbial interactions, the formula (N(N-1)/2) was used where N = number of vertex or ASVs.

### Identification of hydrocarbon degradation proteins in reference genomes

Protein fasta files were derived from annotations of complete genomes from the following candidate novel hydrocarbon-degrading bacteria: *Lacinutrix algicola*, *Lacinutrix himadriensis*, *Lacinutrix jangbogonensis*, *Lacinutrix mariniflava*, *Lacinutrix shetlandiensis*, *Lacinutrix* sp. 5h374, *Lacinutrix* sp. Bg1131, *Lacinutrix* sp C3R15, *Lacinutrix* sp. HL-RS19T, *Lacinutrix venerupis*, *Lacinutrix* sp. Hell90, Candidatus *Pseudothioglobus singularis* PS1 and *Halarcobacter anaerophilus*. Complete genomes for these bacteria were obtained from NCBI Genome database. For searching hydrocarbon degradation proteins, the orthology-based Proteinortho (46) program was used in combination with the Hydrocarbon Aerobic Degradation Enzymes and Genes (HADEG) database (47). The parameters -identity=50 and -conn=0.3 were implemented to reduce the number of false-negative hits and promote the correct annotation of orthologous proteins, as reported elsewhere (47).

## Results

### Microbial biomass and hydrocarbon biodegradation increase with a high nutrient supply

Under oil-amended conditions, high nutrient biostimulation using ORN7a medium resulted in biomass increasing by 100-fold during the 15-week incubation period relative to the ambient and low nutrient conditions (two-way ANOVA test, presence of oil p-value 0.0025, level of nutrients p-value 8.55e-8) (Figure 1A). Declines in n-C_17_/pristane, n-C_18_/Phytane and 3MP/9MP revealed a corresponding increase in crude oil biodegradation in high nutrient microcosms, whereas little change was observed in the lower nutrient systems (Figure 1B). These results show that nutrient addition enhances microbial growth and promotes hydrocarbon degradation by this Arctic Ocean microbiome.

**Figure 1:**
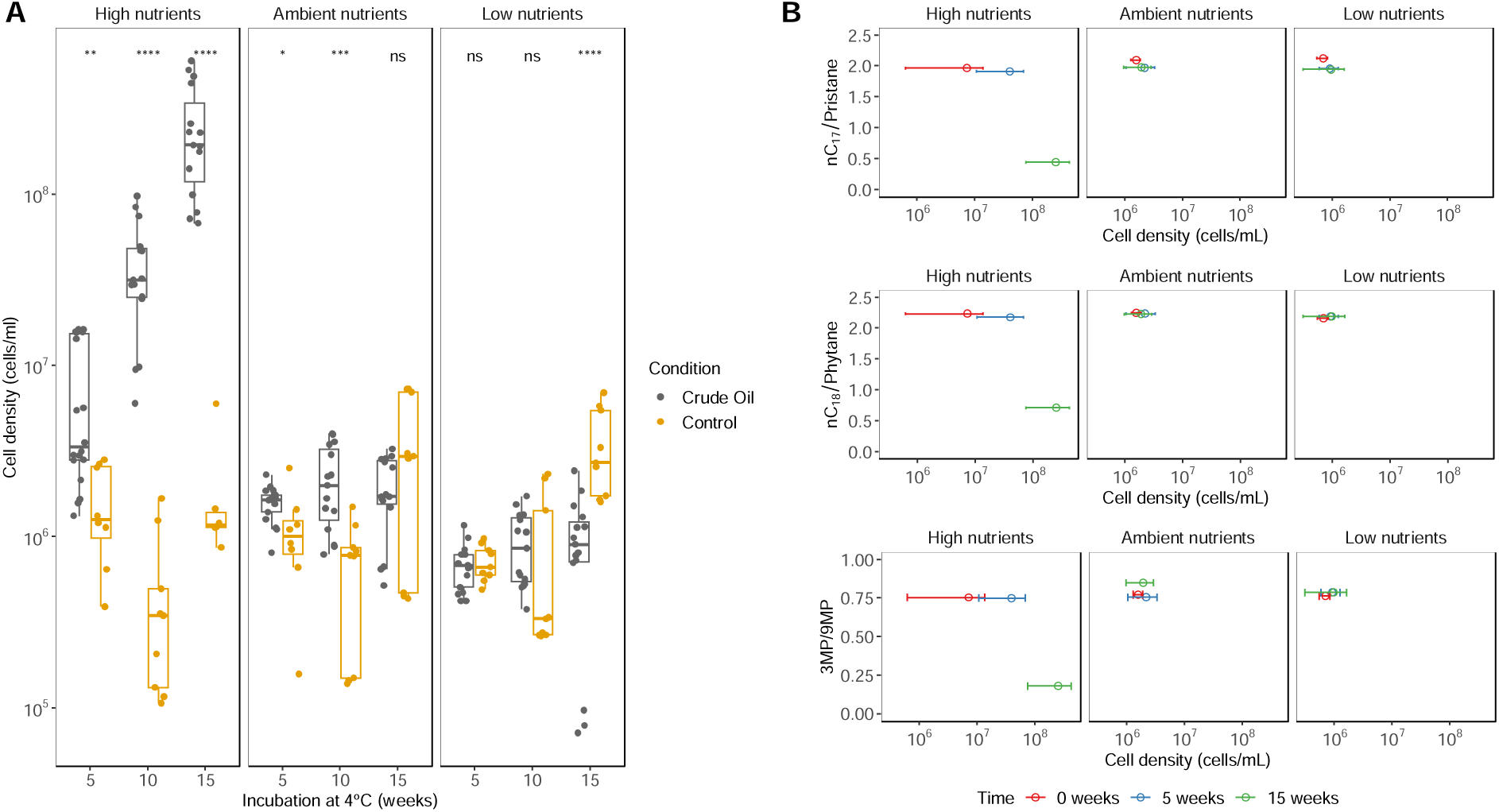
Enhancement of cell density and hydrocarbon degradation in response to nutrient addition. (A) Average values of cell density in each microcosm after 5, 10 and 15 weeks of incubation. Corresponding biodegradation of crude oil hydrocarbons was assessed by tracking ratios of (B) nC_17_/pristane, nC_18_/phytane, and 3-methylphenanthrene/9-methylphenanthrene (3MP/9MP) as a function of changing cell density. Decreases in these hydrocarbon ratios indicate an increasing rate of biodegradation in the presence of high nutrients. Statistical significance: **** : p<= 0.0001, ***: p <=0.001, **: p <=0.01, *: p<=0.05, ns: p> 0.05.

### Microbial diversity response to crude oil contamination and nutrient supplies

Alpha diversity, represented by the number of observed ASVs, decreased significantly over time in microcosms amended with crude oil under high nutrient conditions (two-way ANOVA test, presence of oil p-value 1.27e-11, level of nutrients p-value 3.86e-07) (Figure 2A). In contrast, alpha diversity did not show any significant changes in the presence of crude oil with low or ambient nutrient levels. Microbial community dissimilarity (beta diversity) comparisons revealed that microcosms with the same nutrient supply cluster together (Figure 2B, stress = 0.2472, Figure S1 Shepard plot non-metric R2 = 0.939), with the high nutrient regime differing significantly from the low and ambient nutrient systems (adonis function, Treatment R2 stat 0.09 p-value 9.99e-05: Media R2 stat 0.14707 p-value 9.99e-05). This significant difference is not the consequence of the variance dispersion of the samples within each treatment (betadisper Anova test, Treatment p-value 0.4935: Media p-value 0.8519), by rather driven by microbiome community composition. (Figure 2B).

**Figure 2:**
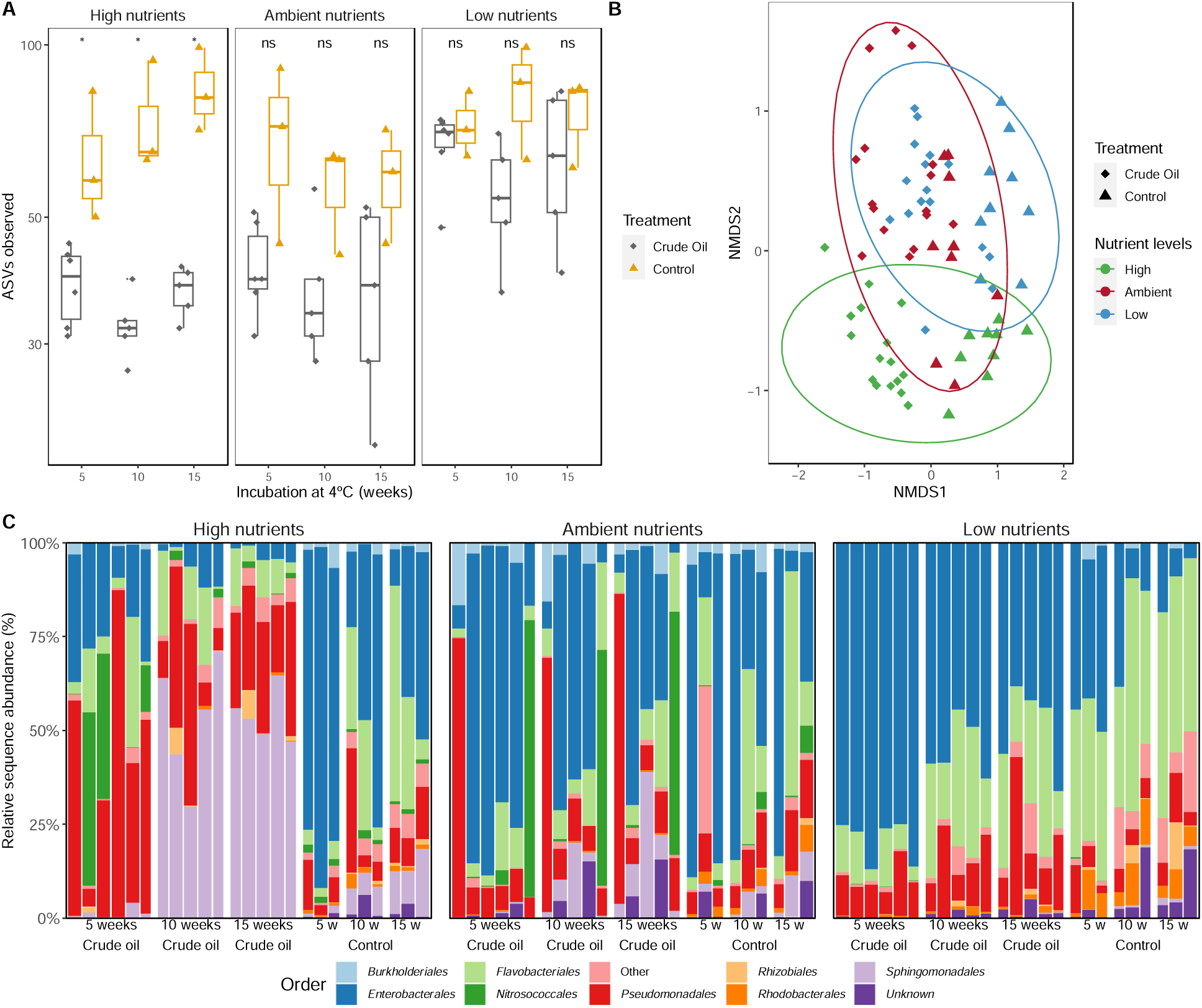
Ecological responses of the Arctic marine microbiome to crude oil contamination and nutrient availability. (A) Alpha diversity values, measured by the number of observed ASVs (richness), in each microcosm after 4°C incubation for 5, 10 and 15 weeks. (B) Non-metric multidimensional scaling (NMDS) of Bray-Curtis dissimilarity of microbial communities exposed to different crude oil and nutrient combinations. (C) Microbial community composition showing top 8 dominant microbial orders in each microcosm grouped by nutrient supply. Each nutrient condition is divided into 6 subgroups, ordered as follows: crude oil-contaminated at 5 weeks of incubation at 4°C (six replicate microcosms), crude oil-contaminated at 10 weeks of incubation at 4°C (five replicate microcosms), crude oil-contaminated at 15 weeks of incubation at 4°C (five replicate microcosms), no-oil control microcosms at 5 weeks of incubation at 4°C (three replicate microcosms), no-oil control microcosms at 10 weeks of incubation at 4°C (three replicate microcosms), and no-oil control microcosms at 15 weeks of incubation at 4°C (three replicate microcosms). Statistical significance: **** : p<= 0.0001, ***: p <=0.001, **: p <=0.01, *: p<=0.05, ns: p> 0.05.

Microbial community shifts were observed in response to nutrient availability and the duration of incubation (Figure 2C) with specific taxa emerging in microcosms with different nutrient treatments. After five weeks of incubation in the presence of crude oil, members of the orders Pseudomonadales (mean relative abundance = 44.85%) and Nitrosococcales (mean relative abundance = 16.22%) were predominant under high nutrient conditions. By week 10, the order Sphingomonadales (mean relative abundance = 52.67%) surpassed Nitrosococcales in abundance (Figure 2C). By comparison, a bloom of Flavobacteriales (mean relative abundance = 19.9%) was observed by week 10 in no-oil control microcosms in the presence of high nutrients.

Under ambient nutrient conditions, crude-oil amended microcosms exhibited an increase in the abundance of Sphingomonadales (mean relative abundance = 11.85%) whereas no-oil controls featured an increase of Flavobacteriales (mean relative abundance = 25.86%) over the 15 weeks of incubation. In the low nutrient microcosms, the microbial community was dominated by Enterobacterales, Flavobacteriales and Pseudomonadales in the presence of crude oil, while Rhodobacterales and Flavobacteriales were dominant without crude oil amendment (Figure 2C). These findings suggest that bacterial taxa belonging to Sphingomonadales are one of the main bacterial taxa involved in crude oil biodegradation in the Arctic waters sampled here, and their activity is enhanced by supplementary nitrogen and phosphorus.

### Differential abundance analysis uncovers both known and novel microbial markers for crude oil biodegradation

Comparing cell density-corrected microbial community structure in oil-contaminated and non-contaminated microcosms using differential abundance analysis (48–50) identified a total of 41 microbial markers at the ASV level associated with crude oil amendment. In microcosms amended with crude oil in the presence of high nutrient concentrations, 22 microbial markers were identified, including bacterial taxa that have been previously described as hydrocarbonoclastic such as *Oleispira*, *Algibacter*, *Shewanella*, *Neptunomonas*, *Halomonas* and *Pseudomonas* (9, 10, 12, 14, 15, 51, 52) (Figure 3A). Microcosms with ambient nutrient conditions featured putative hydrocarbon-degraders *Salegentibacter*, *Zobellia*, *Oceanicoccus* and *Shewanella* (51, 53). (Figure 3B). Under low nutrient conditions, lineages known to include hydrocarbon degraders were enriched, including *Thalassotalea*, *Planococcus*, *Psychromonas* and *Cognaticolwellia* (54, 55).

**Figure 3:**
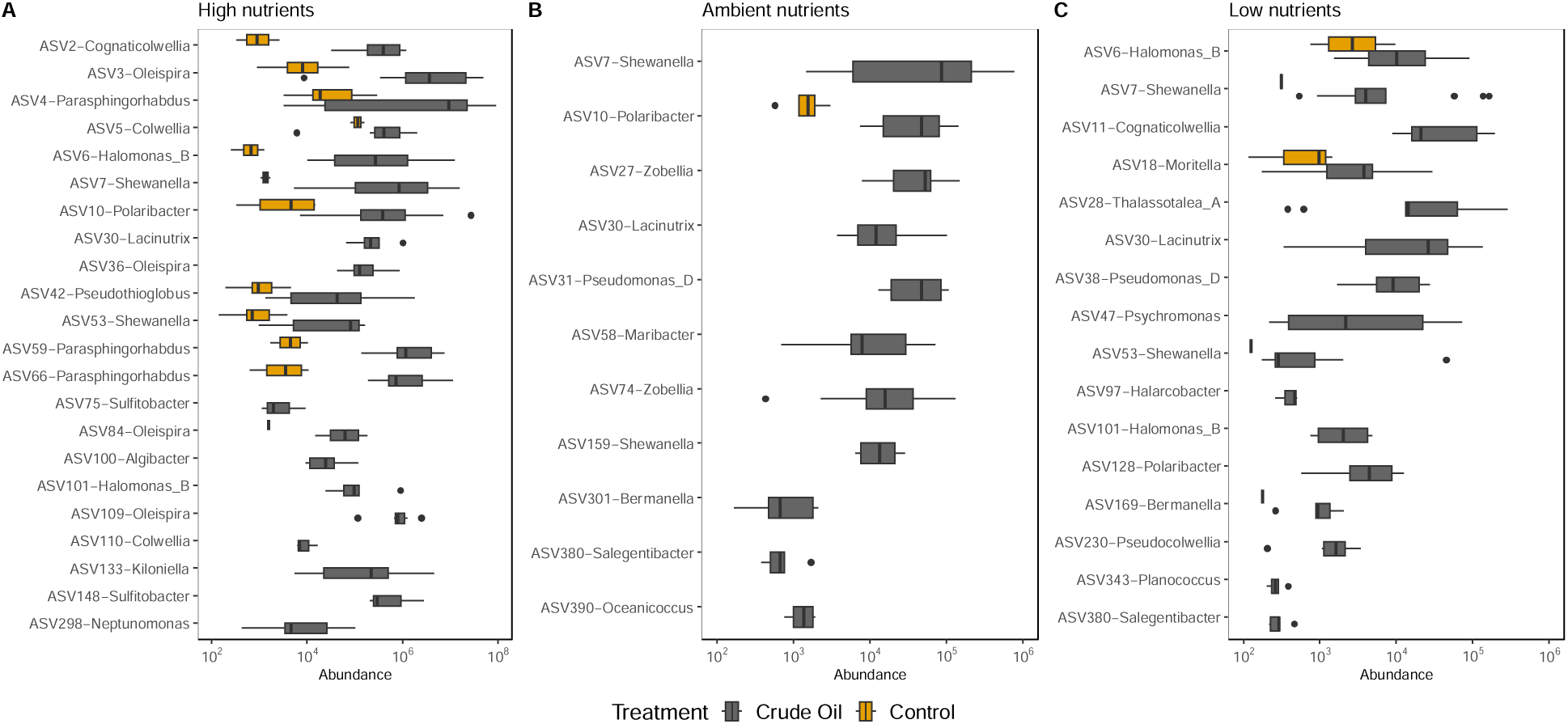
Differential abundance analysis identifies microbial markers for oil-contaminated microcosms. The identification of differentially abundant ASVs under hydrocarbon conditions was performed by comparing the cell count-adjusted total abundance between crude-oil contaminated microcosms (gray) and non-amended microcosms (yellow) using White’s non-parametric test with BH correction in (A) High nutrients, (B) Ambient nutrients, and (C) Low nutrients.

This analysis also highlighted bacterial groups that are not typically associated with crude oil biodegradation. Members of the genus *Lacinutrix, Halarcobacter* and Candidatus *Pseudothioglobus* were consistently associated with crude oil amended across different nutrient regimes (Figure 3).

### Investigating reference genomes of oil-associated genera

Differential abundance analysis revealed the genera *Lacinutrix*, *Halarcobacter* and *Pseudothioglobus* as potential novel microbial markers for oil-contaminated marine environments in the Arctic. To confirm hydrocarbon degradation potential within these genera, orthology-based identification analysis was conducted on reference genomes from Candidatus *Pseudothioglobus singularis* PS1*, Halarcobacter anaerophilus* and 10 different *Lacinutrix* species. Within the genome of *Halarcobacter anaerophilus* various genes associated with the meta-cleavage pathway were identified (56). This genomic pathway is responsible for converting aromatic compounds into intermediates of the TCA cycle (dmpP, dmpN, xylF, xylG, xylH, xylI, xylJ and xylK). All of the *Lacinutrix* genomes contain at least one pca gene (pcaI, pcaF, pcaJ) involved in the ortho-cleavage pathway responsible for biodegradation of aromatic compounds (57, 58). *Lacinutrix* genomes also contain paa genes (paaA, paaB, paaC, paaK) that have been implicated in ring hydroxylation during the aerobic metabolism of phenylacetate and activation of phenylacetate to phenylacetate-CoA (59–61). *Lacinutrix shetlandiensis* specifically harbors the putative toluene biodegradation gene pchA which encodes 4-hydroxybenzaldehyde dehydrogenase (Supplementary Table 1) (62, 63). In the Candidatus *Pseudothioglobus singularis* PS1 genome, one sequence is associated with alkG rubredoxin gene, which is a small essential electron transfer protein in the alkane hydroxylase system (47, 64).

### Genomic traits associated with oil-associated genera

Previous studies have indicated a connection between genomic traits and trophic strategies (18, 65). To explore the potential correlation between genomic traits and trophic strategy in Arctic microbial communities associated with oil contamination studied here, genome size, genomic GC content, and the total number of coding sequences (CDS) among related genomes were inferred (19) for each ASV present in each sample. This involved phylogenetic placement of each ASV on a 16S rRNA gene reference tree created from completed genomes and inferring its genomic traits (19, 40, 66, 67). Resulting values were weighted based on ASV abundance to enable comparison between different nutrient conditions. This analysis predicted a significant increase in the number of coding sequences in members of the high nutrient microbial community over the course of the 15-week incubation (p-value 0.0087), while remaining relatively constant for ambient nutrient community (p-value 0.54) and decreasing slightly for the low nutrient community (p-value 0.017) (Figure 4A). Accordingly, the community mean number of coding sequences under high nutrient conditions is significantly higher than under low nutrient conditions after 15 weeks (p-value 0.007) (Figure 4A; Figure S2A). Inferred genome size did not significantly increase over time in the high nutrient (p-value 0.082) or ambient nutrient (p-value 0.66) conditions, but significantly decreased during the 15-week incubation with low nutrients (p-value 0.017) (Figure 4B). Inference of genomic GC content revealed high GC content enrichment over time for high nutrient (p-value 0.0043) and ambient nutrient (p-value 0.03) conditions, but not for low nutrient conditions (p-value 0.25) (Figure 4C). These effects were especially pronounced at the end of the incubation period (Figure S2C).

**Figure 4.**
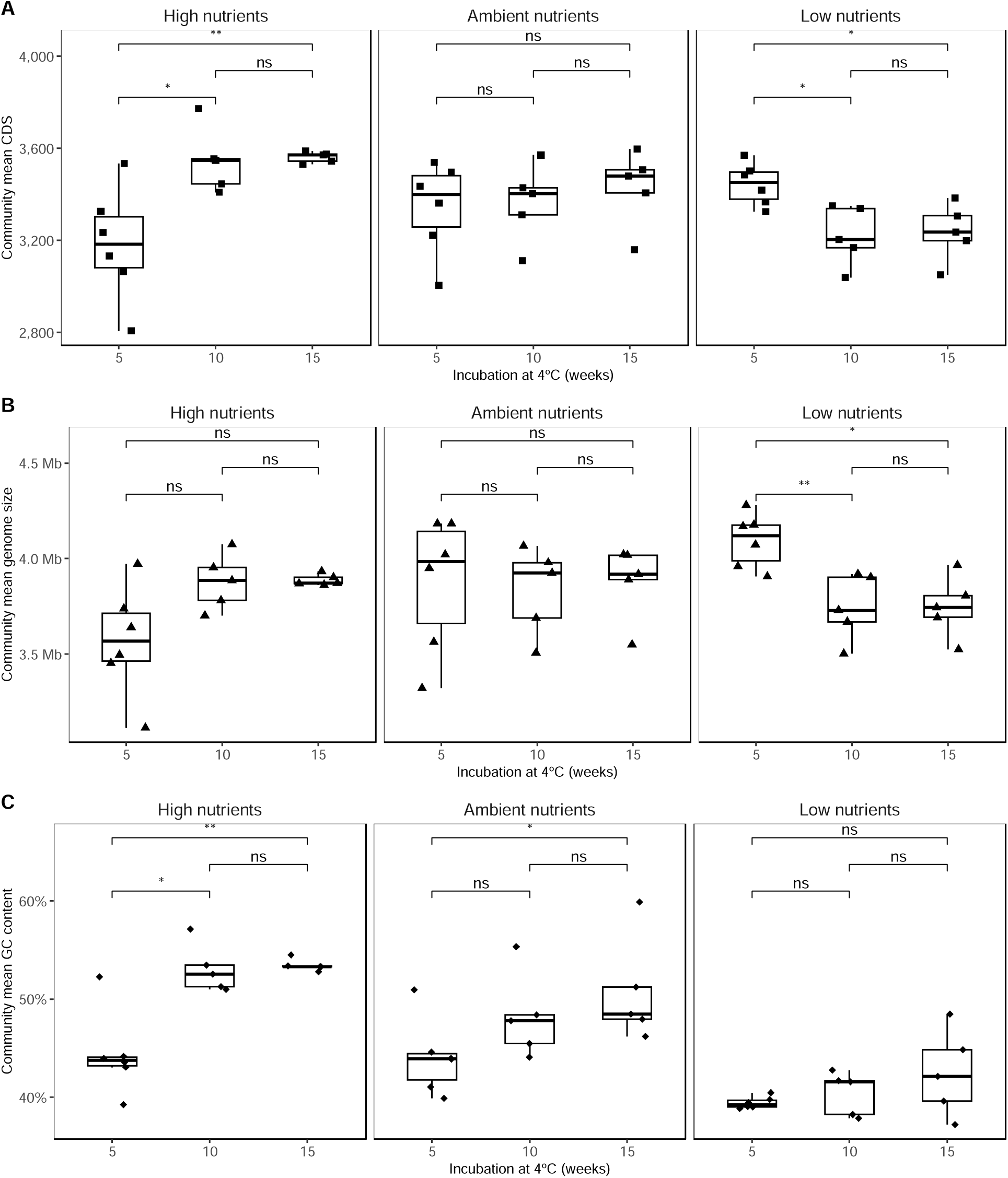
Microbial trophic strategies revealed through inferred community-weighted changes in genomic traits driven by nutrient supply in the presence of crude oil. Community-weight estimate of genomic traits including (A) coding DNA sequences (CDS), (B) genome size and (C) GC content, inferred from 16S rRNA using the metabolic inference pipeline paprica. Statistical analysis was performed comparing different time points using the Wilcoxon Rank test. Statistical significance: ns: p> 0.05, *: p<=0.05, **: p <=0.01, ***: p <=0.001, **** : p<= 0.0001.

### Ecological interactions shift in response to nutrient availability under hydrocarbon exposure

Distinct patterns in microbial interactions based on nutrient conditions were revealed through network analysis in microcosms amended with crude oil. In the presence of high nutrients, 82.9% of microbial taxa exhibited co-occurrence with other ASVs (Figure 5A). The majority of these co-occurrences demonstrate a positive correlation (87.5%), with only small percentage representing negative interactions (12.5%). However, both ambient and low nutrient conditions gave rise to higher bacterial co-occurrence (89.25% and 91.67% respectively) but with the degree of positive co-occurrences diminishing as the nutrient levels became lower (86.48% and 72.72%, respectively) (Figure 5C, Figure S3). At the ASV level, it was observed that certain bacteria showed different patterns of interaction depending on nutrient availability. For instance, *Cognaticolwellia* ASV2 exhibited negative association with *Litorilituus* ASV12 under low nutrient conditions but displayed no association with *Litorilituus* under high nutrient conditions, where it instead exhibits positive co-occurrence with genus *41-12-T18* ASV8 under high nutrient conditions. Similarly, *Polaribacter* ASV10 exhibits a positive co-occurrence with *Methylophilaceae* ASV15 and with *Moritella* ASV18 under ambient nutrient conditions, while no associations are predicted for *Polaribacter* ASV10 under high nutrient conditions. *Lacinutrix* ASV26 exhibits a positive association with bacterial taxa from the *Flavobacteriaceae* family in high nutrient conditions, and a negative association with *Shewanellaceae* taxa under low nutrient conditions.

**Figure 5.**
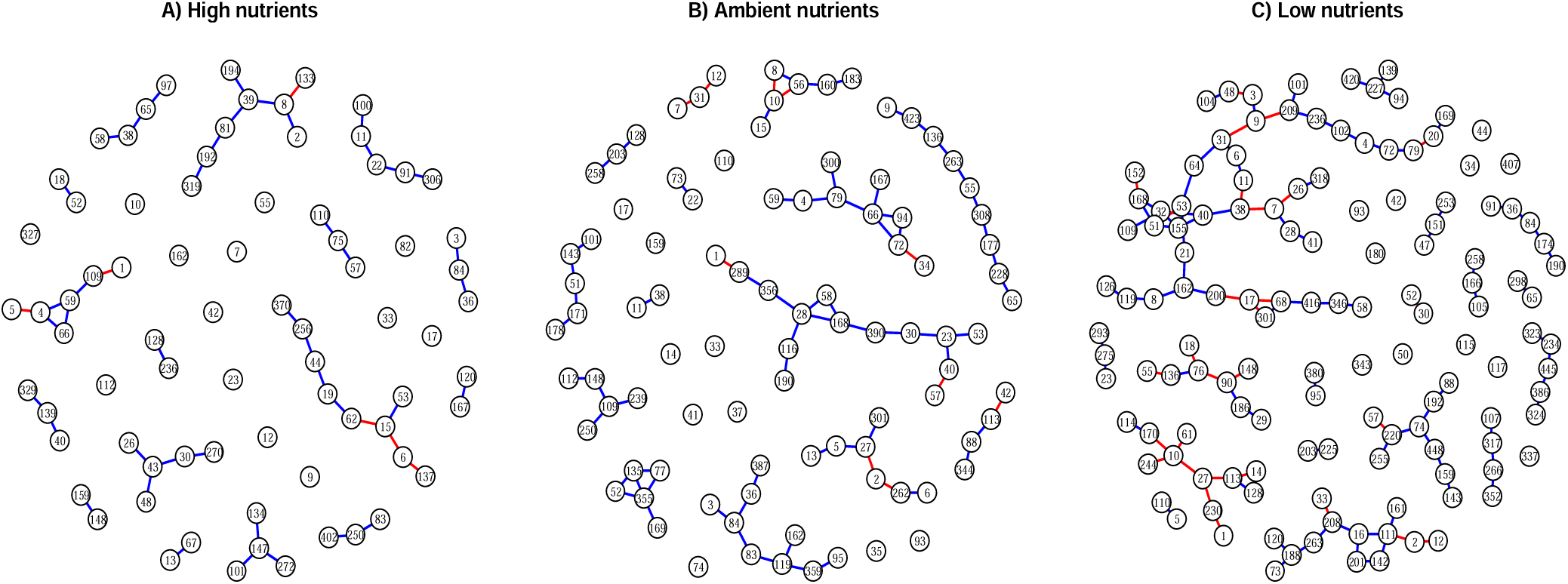
Co-occurrence network analysis in (A) high nutrient, (B) ambient nutrient, and (C) low nutrient microcosms under crude-oil amended conditions. ASVs with a minimum of <2 reads were removed before the analysis. Co-occurrence network analysis was conducted using SPIEC-EASI with the MB method. Each node represents one ASV found in the community, while the colour of the edges indicates positive (blue) or negative (red) co-occurrence associations.

## Discussion

Comparing the diversity in microbial populations using differential abundance analysis of amplicon libraries is valuable for highlighting previously unrecognized and potentially important microbial taxa that might otherwise go unnoticed (49, 50). Combining cell count and 16S rRNA gene diversity data for each microcosm enables total 16S rRNA gene abundances for different groups to be estimated. This eliminates the compositional effect of relative abundance data, which can otherwise increase false positive rates of microbial marker detection (48). In the case of hydrocarbonoclastic taxa, while many microbial lineages known to include hydrocarbon-degrading bacteria were enriched in the experiments performed here (Figure 2C; Figure 3) (9, 12, 51) , the microbiomeMarker pipeline also identified less well-understood genera *Halarcobacter*, Candidatus *Pseudothioglobus* and *Lacinutrix* as being among the most abundant groups in cold Northwest Passage seawater exposed to crude oil.

*Lacinutrix* is a genus within the family *Flavobacteriaceae* comprising 13 gram-negative and strictly aerobic species isolated from marine and polar environments (68, 69). Comparative genomic analysis of these isolates identified genes for cold shock proteins and nitrous oxide reductase, which aid in adapting to cold environmental conditions (70). However, prior studies do not report any ability of *Lacinutrix* spp. related to hydrocarbon biodegradation. Here, examination of 10 *Lacinutrix* genomes, 4 of which were isolated from polar environment (*L. himadriensis, L. jangbogonesis, L. Shetlandiensis* and *L.* sp. Bg11-31), using orthology-based analysis for hydrocarbon biodegradation genes (46, 47) revealed extensive capacity for the biodegradation of aromatic compounds by *Lacinutrix* spp.

Unlike most other groups enriched in crude oil amended microcosms, *Lacinutrix* ASV30 exhibited a prominent response to crude oil addition regardless of whether the nutrient level was high, ambient or low (Figure 3). This aligns with characteristics of facultative oligotrophic bacteria capable of surviving in low-nutrient environments and adapting to higher levels of nutrients when they become available (71–73). This suggests that *Lacinutrix* ASV30 are versatile bacteria that may respond to oil spills in the Arctic marine environment even in the days prior to a biostimulation intervention (ambient nutrients), thereby potentially priming the population to continue or accelerate a bioremediation response after a biostimulation intervention (high nutrients). Other genera known to include facultative oligotrophs identified in this polar environment include *Halomonas* ASV6 and ASV101 and *Shewanella* ASV7 and ASV53 (74), in agreement with ASVs from these genera also being enriched in the presence of oil with varying levels of nutrients (Figure 3). Enrichment of *Lacinutrix* ASV30 in oil-amended microcosms may also be explained by recently described surfactant-like Lyso-Ornithine lipids produced by an Arctic marine *Lacinutrix* isolate (75). Similar to the chemical dispersants used to combat aquatic oil spills (76), these biosurfactant compounds reduce oil-water interfacial tension leading to the emulsification of oil and the formation of smaller oil droplets. The resulting increase in oil water contact surface area has been shown to enhance biodegradation by hydrocarbonoclastic bacteria (77), such as *Shewanella* (14, 78), *Pseudomonas* (12, 79, 80), *Oleispira* (12, 78) and other groups detected in the oil-amended experiments (Figure 3).

*Halarcobacter* was among the microbial groups recently identified in fuel reactors containing PetroF76 and FT-F76, along with other putative hydrocarbon degraders such as *Marinobacter* (81). Candidatus *Pseudothioglobus singularis*, a newly described Gammaproteobacteria, is a group that has been intensively characterized as a representative culture from within the SUP05 clade for its utilization of organic carbon and methylated amines for energy acquisition in marine environments (82). Nevertheless, until now *Halarcobacter* and *Pseudothioglobus* have not been investigated for genes or biosurfactants related to hydrocarbon degradation (81). Genomic inspection revealed here that *Halarcobacter anaerophilus* and Candidatus *Pseudothioglobus singularis* possess alkG rubredoxin and meta-cleavage pathway genes that could potentially be responsible for aromatic compound metabolism, suggesting roles for these populations in crude oil bioremediation in cold Arctic waters. Rubredoxin proteins are essential components in many biochemical processes, including lipid homeostasis and fatty acid beta-oxidation (83). As electron carriers these proteins play broader roles that can be independent of hydrocarbon degradation. Further analysis is necessary to validate the presence of hydrocarbon-degrading alkG rubredoxin genes for *Halarcobacter* and Candidatus *Pseudothioglobus*. Genomic traits such as genome size, GC content, codon usage bias and ribosomal RNA gene copy number are determinants of ecological adaptation (18). High-nutrient environments tend to feature microbial lineages with particular genomic traits indicative of trophic adaptation. Selection for microbial populations with larger genomes in these situations (18), suggests greater metabolic diversity and the potential to metabolise a wider variety of carbon sources as important traits (84). Higher numbers of coding genes allow bacteria with larger genomes to better respond to environmental perturbations (85). High nutrient microcosms in this study also resulted in an increase in the predicted number of coding sequences (Figure 4A) indicative of fertilization with N and P compounds driving the enrichment of copiotroph populations (18).

Conversely, low nutrient conditions resulted in enrichment of ASVs with corresponding genome predictions featuring smaller genome size (Figure 4B), consistent with a strategy employed by oligotrophic bacteria for maintaining genomic stability in relation to low phosphorus availability (86). Genome size is generally correlated with cell size (85). Although cell size was not measured in this study, higher cellular surface area-to-volume ratios have been reported to be an effective nutrient uptake strategy in oligotrophic environments(85, 87). Smaller cell circumferences reduce the costs associated with cell wall and membrane biosynthesis and maintenance, which is optimal in environments with low nutrient availability (85).

GC content of marine bacterial genomes increases with the concentration of inorganic nitrogen (owing to GC pairs utilizing three nitrogen bonds) and has been observed as water depth increases (88). High nutrient microcosms exhibited a significant increase in inferred GC content within the community over time, whereas low nutrient microcosms remained stable in this regard (Figure 4C). GC-rich genes may have an adaptive strategy for increasing bacterial growth rate, suggesting that high genomic GC content, particularly under copiotroph conditions, could be consistent with enhanced microbial growth (89, 90).

Previous studies have described nutrient availability as playing a crucial role in determining microbial interactions at the species level. Depending on the available nutrients in an environment, a pair of species may either compete or collaborate (91, 92). The experiments presented here demonstrate the significant impact of nutrient availability on ecological associations between microbial taxa in the presence of crude-oil (Figure 5). A high degree of antagonistic microbial associations in low-nutrient environments reflect a less cooperative community (16). Previous studies described that under low nutrient supply, microbes tend to reduce the excretion of secondary metabolites, thereby limiting the potential for resource sharing with neighbouring microbes that also rely on such compounds (16, 91). In agreement with this, lower nutrient availability did not promote a greater proportion of positive associations in Arctic microbial communities (Figure 5). In contrast, it has been reported that in high nutrient conditions, microbes can alter the chemical environment by producing harmful metabolites like antimicrobials or by altering external pH to inhibit competitors, thereby decreasing microbial diversity and positive interactions between taxa (16, 93). Overall, the Arctic marine microbial community at this specific sampling location exhibits positive or negative associations with other taxa depending on nutrient availability.

## Conclusions

This study provides evidence that the availability of nutrients strongly impacts marine microbial ecology and the biodegradation response to crude oil contamination in Arctic waters. These changes in the community are characterized by a reduction in microbial diversity, significant shifts in composition, modification in prevailing genomic traits, and changes in the degree of microbial associations. Differential abundance of ASVs affiliated with *Lacinutrix*, *Halarcobacter* and Candidatus *Pseudothioglobus* alongside other taxa suggests these groups are previously unrecognized markers for hydrocarbon degradation in Arctic marine environments, and furthermore exhibit an ability to respond to crude oil inputs regardless of prevailing nutrient concentrations. These findings have important implications for developing bioremediation strategies to address oil spills in vulnerable Arctic waters.

## Acknowledgments

The authors thank the Canadian High Arctic Research Station for assistance during fieldwork, as well as Rhonda Clark for organizing field work and research administration support. We appreciate assistance from Jianwei Chen on gas chromatography and Carmen Li for performing Illumina MiSeq 16S rRNA gene sequencing. We thank Kim Nightingale and the PRG Laboratory at the University of Calgary for performing GC-MS on hydrocarbon extracts, and Jayne Rattray for assistance with GC-MS data analysis. The authors also gratefully acknowledge Srijak Bhatnagar, Whitney England, and other members of the GEMM research group for their valuable discussions and input.

We declare that are no conflicts of interest.

## Data availability

All raw 16S rRNA gene sequence reads generated for this study have been submitted to the GenBank database under BioProject PRJNA1088749.

