## Supplementary Figures for "Novel oil-associated bacteria in Arctic seawater exposed to different nutrient biostimulation regimes"

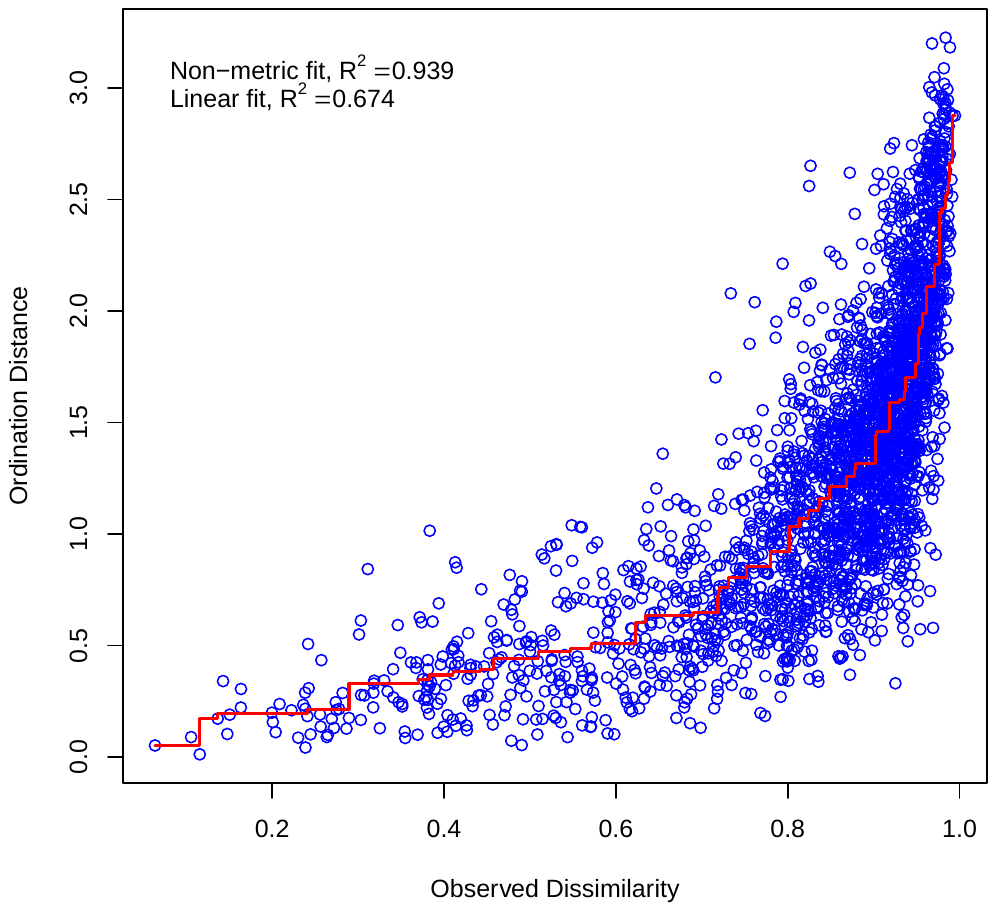

Figure S1. Shepard plot for nonmetric multidimensional scaling identified a strong correlation (R2 = 0.939) between calculated and ordination distances. The proximity of the blue dots to the line reflects how accurately the NMDS solution represents the observed dissimilarity, with closer dots indicating a better fit between NMDS distances and observed dissimilarities.

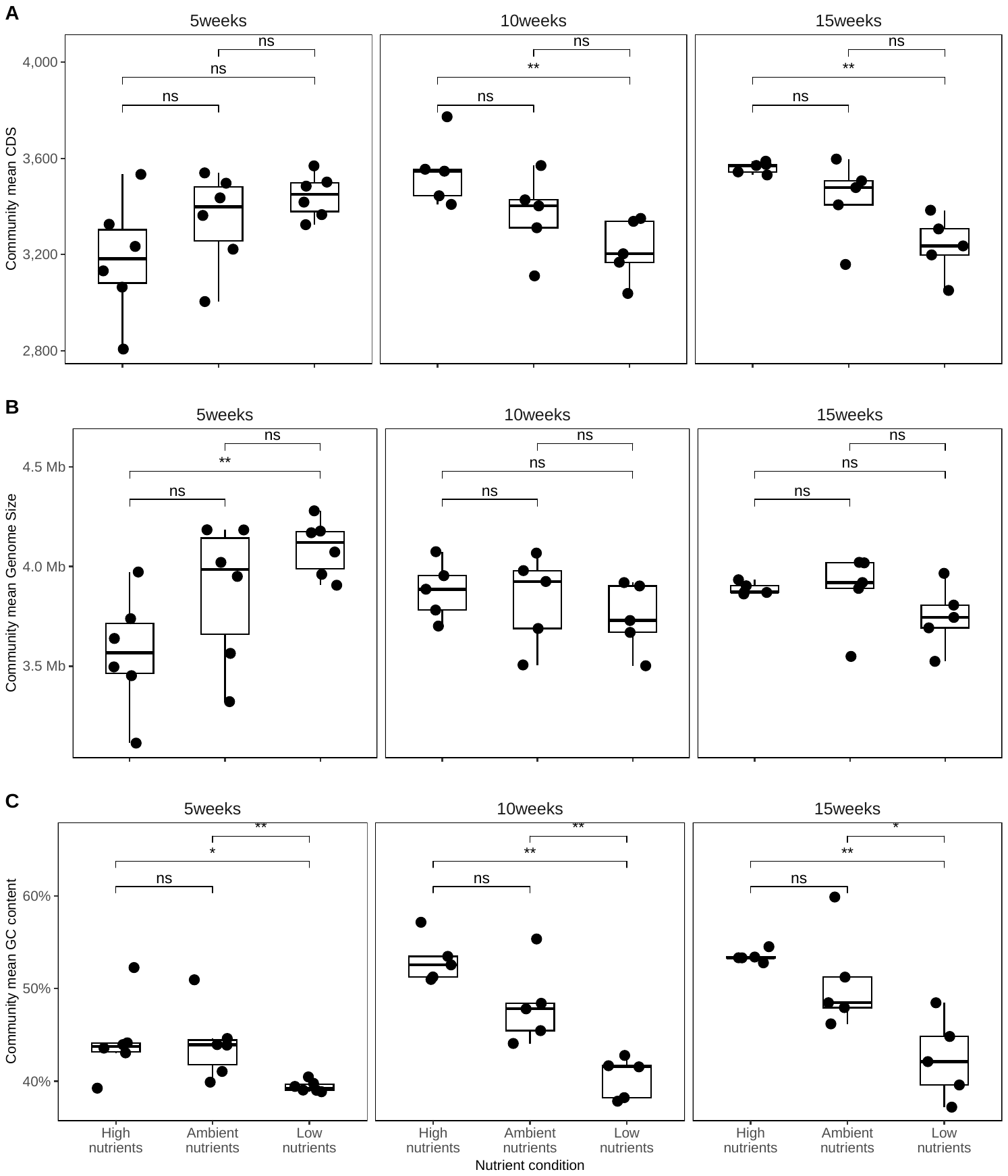

*Figure* *S2.*  Comparison of inferred genomic traits at different time points. Statistical significance: ns: p> 0.05, *: p<=0.05, **: p <=0.01, ***: p <=0.001, **** : p<= 0.0001.

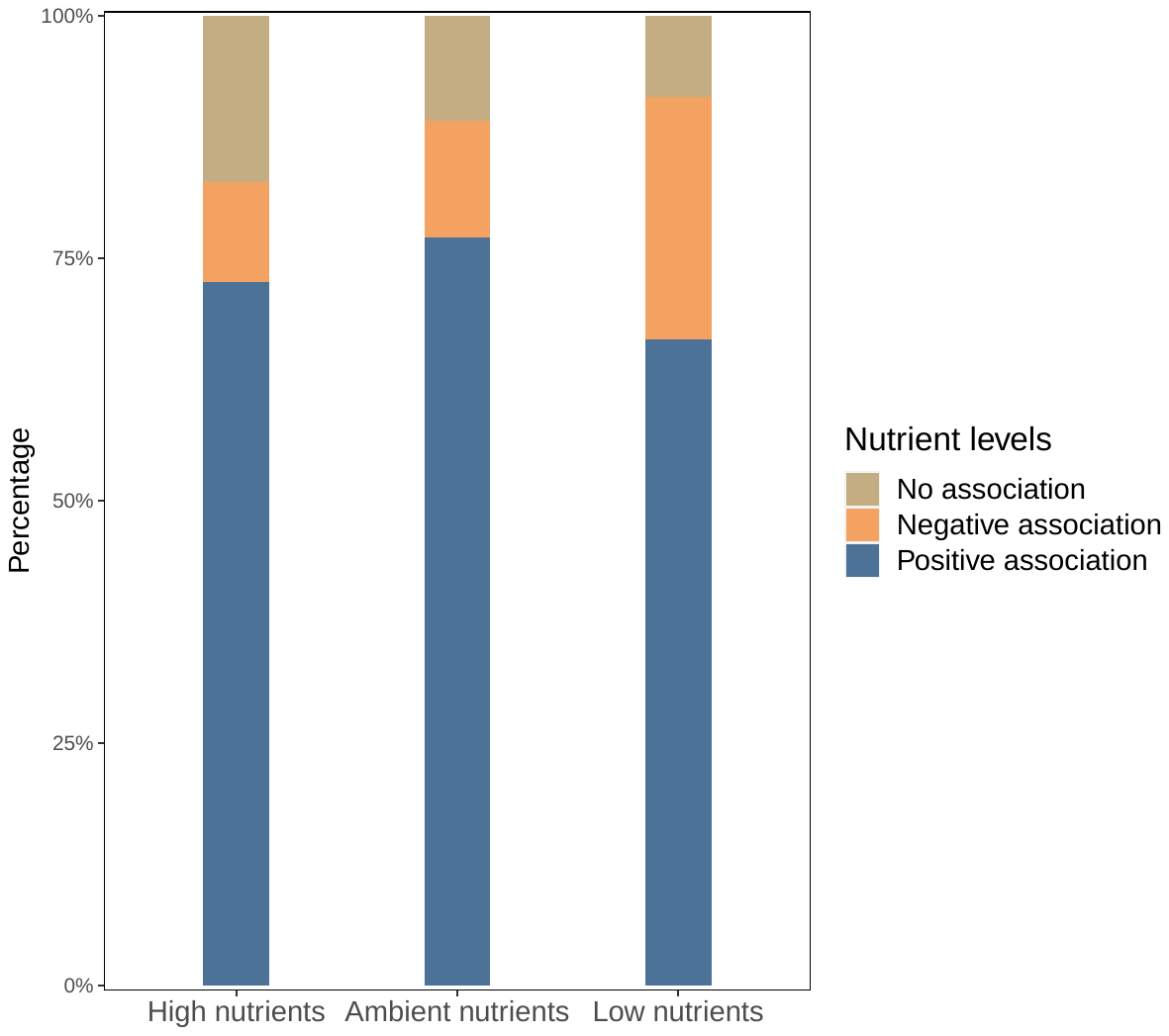

Figure S3. Types of microbial associations of each nutrient condition. Negative microbial associations tend to increase as the nutrient conditions are reduced.
